# Nanoparticles alter the nature and strength of intraploidy and interploidy interactions in plants

**DOI:** 10.1101/2024.03.11.584520

**Authors:** Elizabeth A. Esser, Jiaqi Tan, Na Wei

## Abstract

Engineered nanoparticles have profound impacts on organisms, yet there is limited understanding of how nanoparticle exposure shapes species interactions that are key for natural community dynamics. By growing plants of the same (intraploidy) and different ploidy levels (interploidy) of *Fragaria* in axenic microcosms, we examined the influence of nanoparticles on species interactions in polyploid and diploid plants. We found that, under copper oxide (CuO) nanoparticle exposure, polyploids experienced reduced competition when growing with polyploids (the effect of polyploids on polyploids, RII_8*x*,8*x*_), and a shift towards facilitation when growing with diploids (the effect of diploids on polyploids, RII_8*x*,2*x*_). This reduction in competitive interactions in polyploids, in line with the stress gradient hypothesis, was primarily caused by nanoscale effects, because the strength of competitive interactions (RII_8*x*,8*x*_ and RII_8*x*,2*x*_) remained relatively unchanged under CuO bulk particles compared to control conditions. In contrast, diploids experienced a shift from facilitation (RII_2*x*,2*x*_ and RII_2*x*,8*x*_) under control conditions to neutrality under CuO nanoparticles, with a similar reduction in facilitation observed with both nanoparticles and bulk particles. These findings underscore ploidy specific interaction dynamics and the need of considering species interactions when predicting organismal responses to nanoparticle pollution in ecological communities.

## 1. Introduction

Polyploidy, or whole-genome duplication (with or without hybridization), is widespread in flowering plants and has occurred at least once in the evolutionary history of all angiosperms (Jiao et al., 2011; Vanneste et al., 2014; Cai et al., 2019; One Thousand Plant Transcriptomes Initiative, 2019; Wu et al., 2020). Polyploid plants often diverge from diploid counterparts in phenotypic characteristics, including changes in morphological and physiological traits that may influence organismal fitness (Maherali et al., 2009; Wei et al., 2019; Wei et al., 2020; Guo et al., 2023). Such phenotypic divergence is often thought to arise from the increase in genome size and chromosome copies, which can fuel phenotypic and functional innovations as well as genomic diversification and versatility (Levin, 1983; Comai, 2005; Ramsey and Ramsey, 2014; Soltis et al., 2016; Bomblies, 2020; Van de Peer et al., 2021). As such, polyploids are hypothesized to have an increased ability to withstand abiotic stress and biotic antagonism (e.g., competitive interactions) (Stebbins, 1971; Levin, 1983; Ramsey and Ramsey, 2014; Cai et al., 2019; Wu et al., 2020; Van de Peer et al., 2021). While recent studies have shown evidence for a polyploid advantage in competitive interactions in a handful of polyploid complex systems (Maceira et al., 1993; Collins et al., 2011; Thompson et al., 2015; Rey et al., 2017; Čertner et al., 2019; Cheng et al., 2020; Guo et al., 2023), notable exceptions exist (Thompson et al., 2015; Anneberg et al., 2023). It remains largely unknown whether the hypothesized polyploid advantage holds across various environmental conditions, particularly in the novel environments of the Anthropocene. These novel environmental conditions, including exposure to new pollutants like engineered nanoparticless, are likely to alter the nature and strength of species interactions (Peng et al., 2017; Wu et al., 2019; Menicagli et al., 2023; Zuo et al., 2024). Therefore, understanding species interactions in these novel environments is essential for predicting the future interactive dynamics of polyploid and diploid plants.

One of the emerging environmental threats in the Anthropocene arises from engineered nanoparticles (Colvin, 2003; Peralta-Videa et al., 2011; Aslani et al., 2014; Zhang et al., 2019; Ameen et al., 2021). Nanoparticles, with unique properties such as nanoscale size (1–100 nm), large relative surface area, novel physiochemical attributes, and high reactivity, have revolutionized various industries (e.g., electronics, cosmetics, medicine, and agriculture), leading to an annual global production of millions of metric tons (Biswas and Wu, 2005; Zhang et al., 2019; Ameen et al., 2021; Martinez et al., 2021; Keller et al., 2023). Through production, use, disposal, runoff, and atmospheric deposition, thousands of metric tons of engineered nanoparticles are discharged into terrestrial and aquatic ecosystems each year (Keller et al., 2023). This increasing accumulation poses a threat to ecosystem functioning by adversely affecting plants, microbes, and animals (Peralta-Videa et al., 2011; Aslani et al., 2014; Ameen et al., 2021; Khan et al., 2021; Gakis et al., 2023; Chen et al., 2024). The negative impacts on organisms can arise through various mechanisms, such as nanoscale size facilitating entry into cells, interference with cell division, DNA damage, and induction of cell death (Singh et al., 2009; Khan et al., 2021; Yang et al., 2021). Aggregation on cell surfaces can disrupt nutrient transport (Ameh and Sayes, 2019; Yang et al., 2021), while inherent material characteristics, particularly heavy metals, can lead to the creation of harmful reactive oxygen species and biomolecule oxidation (Singh et al., 2009; Ameh and Sayes, 2019; Singh et al., 2019; Khan et al., 2021). Notably, toxicity levels may differ between bulk and nano forms of a material. If the toxic effects are mainly attributed to material effects, organisms exposed to both bulk and nano forms may exhibit similar levels of toxicity. However, if toxicities primarily result from nanoscale effects, organisms exposed to nanoparticles are expected to experience stronger toxicity compared to those exposed to the bulk counterparts. For example, copper oxide (CuO) nanoparticles have been shown to exhibit toxicity to plants and animals at doses ten times lower than those required for CuO bulk particles (Ameh and Sayes, 2019). Given their common application in agricultural and environmental remediation as fertilizers and pesticides, CuO nanoparticles have been extensively investigated in a variety of key crops. These include, for example, rice (*Oryza sativa*), maize (*Zea mays*), wheat (*Triticum aestivum*), soybean (*Glycine max*), cotton (*Gossypium hirsutum*), spinach (*Spinacia oleracea*), zucchini (*Cucurbita pepo*), sweet potato (*Ipomoea batatas*), and alfalfa (*Medicago sativa*) (Hong et al., 2015; Le Van et al., 2016; Singh and Kumar, 2016; Bradfield et al., 2017; Gao et al., 2018; Yusefi-Tanha et al., 2020; Roy et al., 2022). While these previous studies have revealed adverse effects of nanoparticles on individual organisms, such as reduced plant germination, nutrient uptake, and biomass (Rajput et al., 2017), it remains largely unknown how nanoparticle exposure influences species interactions (Peng et al., 2017; Wu et al., 2019; Menicagli et al., 2023; Zuo et al., 2024), especially in wild plants.

While polyploids have been hypothesized to exhibit competitive dominance over diploids (Wei et al., 2019; Guo et al., 2023), the impact of nanoparticles on the competitive interactions between polyploids and diploids remains unclear. The dynamics of plant–plant interactions, in terms of both nature and magnitude, can vary under different environmental conditions (Bertness and Callaway, 1994; Maestre et al., 2009; He et al., 2013; Zhang and Tielborger, 2020). According to the stress gradient hypothesis (SGH), the magnitude of competition may decrease and potentially shift towards facilitation in more stressful environments (Bertness and Callaway, 1994; Maestre et al., 2009). SGH also suggests that the type of stress experienced can influence species interactions (He et al., 2013). While increased resource stress (e.g., water or nutrient deficiency) often reduces the magnitude of competition, increased non-resource stress (e.g., cold, heat, salinity) may lead to shifts from competition to facilitation (He et al., 2013; Zhang and Tielborger, 2020). In this study, we aim to investigate how CuO nanoparticles influence the interactions between polyploids and diploids. Owing to, for example, the genomic and phenotypic novelty and versatility of polyploidy (Wei et al., 2019; Guo et al., 2023), polyploids are hypothesized to be competitively dominant (Fig. 1). As a result, in polyploids, the competitive effect of diploids on polyploids (RII_8*x,*2*x*_, 2*x* and 8*x* refer to diploids and octoploids, respectively) is expected to be weaker than the effect of polyploids on polyploids (RII_8*x,*8*x*_ > RII_8*x,*2*x*_). Conversely, in diploids, the competitive effect of polyploids on diploids (RII_2*x,*8*x*_) is expected to be stronger than the effect of diploids on diploids (RII_2*x,*8*x*_ > RII_2*x,*2*x*_). Stress induced by nanoscale effects is expected to reduce intraploidy competition (RII_8*x,*8*x*_ and RII_2*x,*2*x*_) and interploidy competition (RII_8*x,*2*x*_ and RII_2*x,*8*x*_), as opposed to exposure to bulk particles and control conditions (in the absence of the two forms of the material; Fig. 1a,b). However, if stress is attributed to material effects, rather than nanoscale effects, the reduction in competition is expected to be similar between exposure to nanoparticles and bulk particles (Fig. 1c,d). In the absence of stress from nanoparticles or bulk particles, competition is expected to be similar across conditions (Fig. 1e,f). On the other hand, as nanoparticles, such as CuO nanoparticles, are often used as fertilizers (Lowry et al., 2019), competition may be increased due to factors such as increased plant sizes (Fig. 1g,h). However, if polyploidy leads to, for example, genomic instability and slower metabolism (Levin, 1983; Comai, 2005), polyploids may be competitively inferior (diploid dominance: RII_8*x,*2*x*_ > RII_8*x,*8*x*_ and RII_2*x,*8*x*_ < RII_2*x,*2*x*_; Fig. S1) or competitively similar to diploids (competitive symmetry: RII_8*x,*8*x*_ = RII_8*x,*2*x*_ and RII_2*x,*8*x*_ = RII_2*x,*2*x*_; Fig. S2). Their competitive interactions may be weakened due to nanoscale effects or material effects, or strengthened due to fertilization effects (Figs. S1 and S2).

**Fig. 1.**
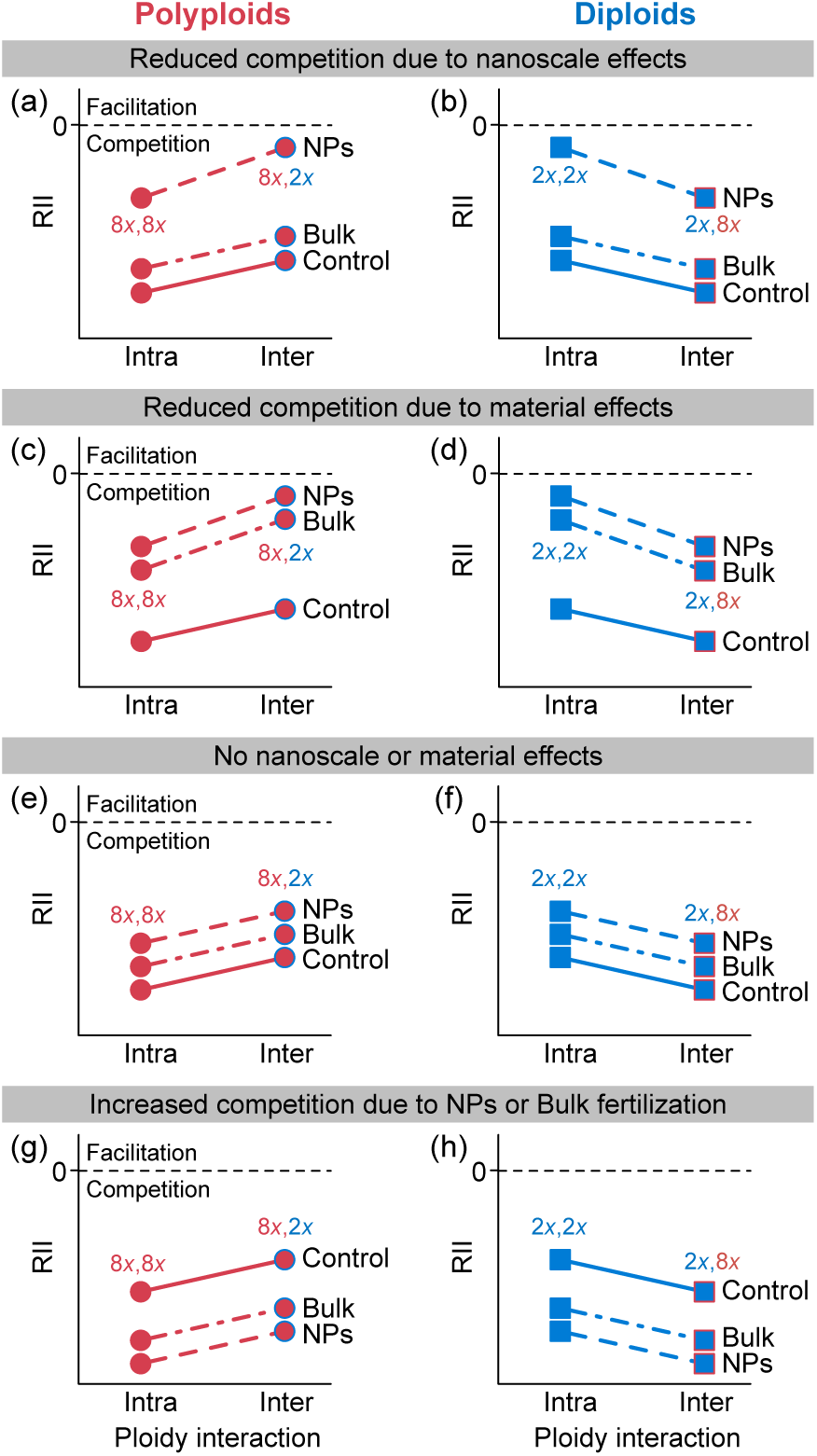
Hypotheses of competitive interactions between polyploids and diploids in response to nanoparticles. (a-f) Plant–plant interactions between ploidy levels are quantified using the relative interaction index (RII), where RII < 0 indicates competition and RII > 0 indicates facilitation. If polyploids (8*x*, octoploid as an example) are stronger competitors, the competitive effect of polyploids on polyploids (RII_8*x*,8*x*_) is expected to be stronger than the effect of diploids on polyploids (RII_8*x*,2*x*_), resulting in stronger intraploidy than interploidy competitions, RII_8*x*,8*x*_ > RII_8*x*,2*x*_. In contrast, diploids are expected to be more affected by interploidy competition (RII_2*x*,8*x*_) than intraploidy competition (RII_2*x*,2*x*_), leading to RII_2*x*,8*x*_ > RII_2*x*,2*x*_. The intraploidy and interploidy competitions may, nevertheless, change under non-resource stress (e.g., nanoparticles and bulk particles of the material). The stress gradient hypothesis (SGH) predicts reduced competition with increased stress. (a, b) If the stress is caused by nanoscale effects rather than material effects, a reduction in competition is expected under nanoparticles (NPs), in contrast to bulk particles (Bulk) and the control (Control) where no nanoparticles or bulk particles are introduced. (c, d) If the stress is caused by material effects, a reduction in competition is expected to be similar under both nanoparticles and bulk particles, relative to the control. (e, f) However, competition may remain unchanged if nanoparticles and bulk particles are not the primary sources of stress that influences plant–plant interactions. (g, h) On the other hand, if nanoparticles and bulk particles provide fertilization, intraploidy and interploidy competitions may increase due to reasons such as increased plant sizes.

**Fig. 2.**
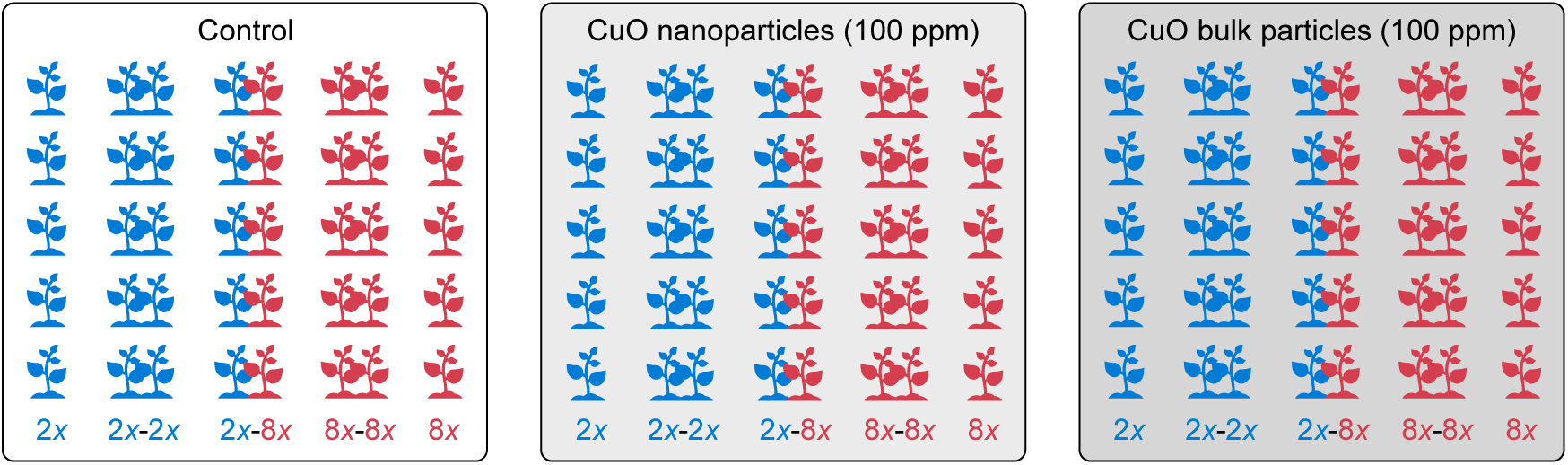
Summary of experimental design. The basic unit of the competition experiment consists of five plant combinations: diploid growing alone (2*x*), diploid growing with intraploidy level (2*x*– 2*x*), diploid growing with polyploid (2*x*–8*x*), polyploid growing with intraploidy level (8*x*–8*x*), and polyploid growing alone (8*x*). This basic unit was replicated five times for each of the three treatments: the control (no copper oxide nanoparticles or bulk particles), copper oxide (CuO) nanoparticles, and CuO bulk particles.

To evaluate the influence of CuO nanoparticles, one of the widely used nanoparticles (Keller et al., 2023), on interactions between polyploids and diploids, we competed diploid and polyploid plants with intraploidy or interploidy individuals under exposure to CuO nanoparticles and bulk particles in axenic microcosms. We aimed to test the outlined hypotheses (Fig. 1; Figs. S1 and S2), by addressing three questions: (1) Do polyploids exhibit competitive dominance over diploids? (2) Does the impact of nanoparticle exposure on intraploidy and interploidy interactions align with predictions from the stress gradient hypothesis? (3) What mechanisms underlie the impact of nanoparticles on species interactions, specifically examining nanoscale effects versus material effects?

## 2. Material and methods

### 2.1. Experimental Design

To examine the intraploidy and interploidy interactions, we focused on diploid (2*x*) *Fragaria viridis* and octoploid (8*x*) *Fragaria virginiana*, two closely related wild strawberry species (Dillenberger et al., 2018; Wei et al., 2019; Liston et al., 2020). Aseptic propagation of the plants was conducted in Murashige and Skoog Basal medium with vitamins and sucrose (PhytoTech Labs, Lenexa, Kansas) within glass culture tubes (25 mm × 150 mm) before the start of the competition experiment. The basic unit of the competition experiment consisted of five plant combinations (Fig. 2): diploid growing alone (2*x*), diploid growing with intraploidy level (2*x*– 2*x*), diploid and polyploid growing together (2*x*–8*x*), polyploid growing with intraploidy level (8*x*–8*x*), and polyploid growing alone (8*x*). The basic units of the competition experiment were subject to stress treatments (CuO nanoparticles, ‘NPs’; CuO bulk particles, ‘Bulk’; and control), with five replicates each treatment: 5 plant combinations × 3 stress treatments × 5 replicates, resulting in 75 total experimental microcosms.

Before the start of the experiment, individual microcosms (55 mL glass tubes, 25 mm × 150 mm) were filled with 4 g Sunshine Redi-Earth Plug & Seedling Potting Mix (Sun Gro Horticulture, Agawam, Massachusetts) that comprises peat moss, vermiculite, dolomitic limestone, and 4 mL of deionized (DI) water. These microcosms were autoclaved three times for 45 min each time. Then CuO nanoparticles (particle size <50 nm; Sigma Aldrich, St. Louis, Missouri) or CuO bulk particles (particle size ~2000 nm, Alfa Aesar; VWR International, Radnor, Pennsylvania), suspended in 1 mL of autoclaved DI water, was added into the NPs treatment (resulting in a final concentration of 100 ppm in microcosms, i.e., 100 mg kg^-1^) and the Bulk treatment (100 ppm), respectively. While exceeding the naturally occurring copper levels in soil (2–50 mg Cu per kg dry weight; Oorts, 2013), this concentration remains lower than those observed in copper polluted soils (Poggere et al., 2023). Additionally, it is comparable to concentrations used in nano-phytotoxicity studies of CuO and other metal (and oxide) nanoparticles (Khan et al., 2021; Gakis et al., 2023). For the control treatment, 1 mL of autoclaved DI water was added. Following the preparation of microcosms, diploid and polyploid plants were transferred into individual microcosms under a sterile laminar flow hood. The microcosms were capped, and plants were grown under a light intensity of 50 μmol m^-2^ s^-1^ for 16 hours a day at 24 °C. To minimize the potential influence of microenvironmental variation, microcosms were randomized daily. Because microcosms in axenic microcosms without additional added water could experience increased evaporation over time, we ended the experiment after three weeks to prevent potential stress confounding due to water loss. In this study, we measured the height of individual plants in the microcosms at both the beginning and end of the experiment.

### 2.2. Statistical Analyses

We evaluated the nature and intensity of plant–plant interactions using the relative interaction index (RII; Armas et al., 2004) that ranges between −1 and 1. A negative RII indicates competition (with more negative values reflecting stronger competition), while a positive RII indicates facilitation (with higher positive values reflecting stronger facilitation). The RII compares plant growth with competitors (B_w_) and without competitors (B_o_): RII = (B_w_ – B_o_)/(B_w_ + B_o_). In this study, we assessed plant growth using plant height as a proxy, given that plant growth in the microcosms was primarily vertical. In addition, we opted for this non-destructive measurement rather than biomass harvesting to minimize the handling of heavy metal and nanoparticles in the microcosms. For each ploidy combination (2*x*–2*x*, 2*x*–8*x*, 8*x*–8*x*; Fig. 2), B_w_ represented the plant height of each diploid or polyploid plant growing with intraploidy or interploidy competitors in individual microcosms. B_o_ was the plant height of the diploid or polyploid growing alone, averaged across replicated microcosms in each treatment (Fig. 2).

To evaluate the hypothesized intraploidy and interploidy competitive effects (Fig. 1), we conducted general linear models (LMs) with RII as the response variable using R v4.2.2 (R Core Team, 2022). The predictors included competition type (intraploidy vs. interploidy), treatment (control, NPs, and Bulk), and their interactions, with initial plant height as a covariate. We did not consider the random effect of microcosm ID, as its impact was not significant, and its inclusion affected model fitting. The LMs were carried out separately for polyploids (8*x*–8*x*, 2*x*– 8*x*) and diploids (2*x*–2*x*, 2*x*–8*x*). Statistical significance (type III sums of squares), least-squares means (LS means) of predictors, and planned contrasts of intraploidy vs. interploidy interactions and treatment effects within the LMs were assessed using packages emmeans (Lenth, 2023) and phia (De Rosario-Martinez, 2023).

## 3. Results

In polyploids, consistent with the stress gradient hypothesis (Fig. 1 and Fig. S3), the competitive interaction between polyploids (RII_8*x*,8*x*_) was significantly lowered under the NPs treatment relative to the control (LS mean contrast: NPs, RII_8*x*,8*x*_ = −0.04; control, RII_8*x*,8*x*_ = −0.16; *F* = 4.1, df = 1, *p* = 0.050; Fig. 3a; Table S1). Such reduction in the intraploidy competition of polyploids was caused by nanoscale effects (Fig. 1a) rather than material effects (Fig. 1c), because unlike the NPs treatment, the Bulk treatment did not lower RII_8*x*,8*x*_ compared to the control (LS mean contrast: Bulk, RII_8*x*,8*x*_ = −0.23; control, RII_8*x*,8*x*_ = −0.16; *F* = 1.4, df = 1, *p* = 0.2). In addition, we found significantly lowered RII_8*x*,8*x*_ under the NPs treatment compared to the Bulk treatment (*F* =10.4, df = 1, *p* = 0.003). Similar to the intraploidy competition of polyploids (RII_8*x*,8*x*_), the competitive effect of diploids on polyploids (RII_8*x*,2*x*_) was also lowered under the NPs treatment (Fig. 3a), especially compared to the Bulk treatment (LS mean contrast: NPs, RII_8*x*,2*x*_ *=* 0.06; Bulk, RII_8*x*,2*x*_ *=* −0.16; *F* = 6.8, df = 1, *p* = 0.013). While the reduction in RII_8*x*,2*x*_ was not significant relative to the control (LS mean contrast: NPs, RII_8*x*,2*x*_ *=* 0.06; control, RII_8*x*,2*x*_ *=* −0.08; *F* = 2.8, df = 1, *p* = 0.1), the nature of the interaction (RII_8*x*,2*x*_) shifted from competition under the control and Bulk treatment to facilitation under the NPs treatment (Fig. 3a).

**Fig. 3.**
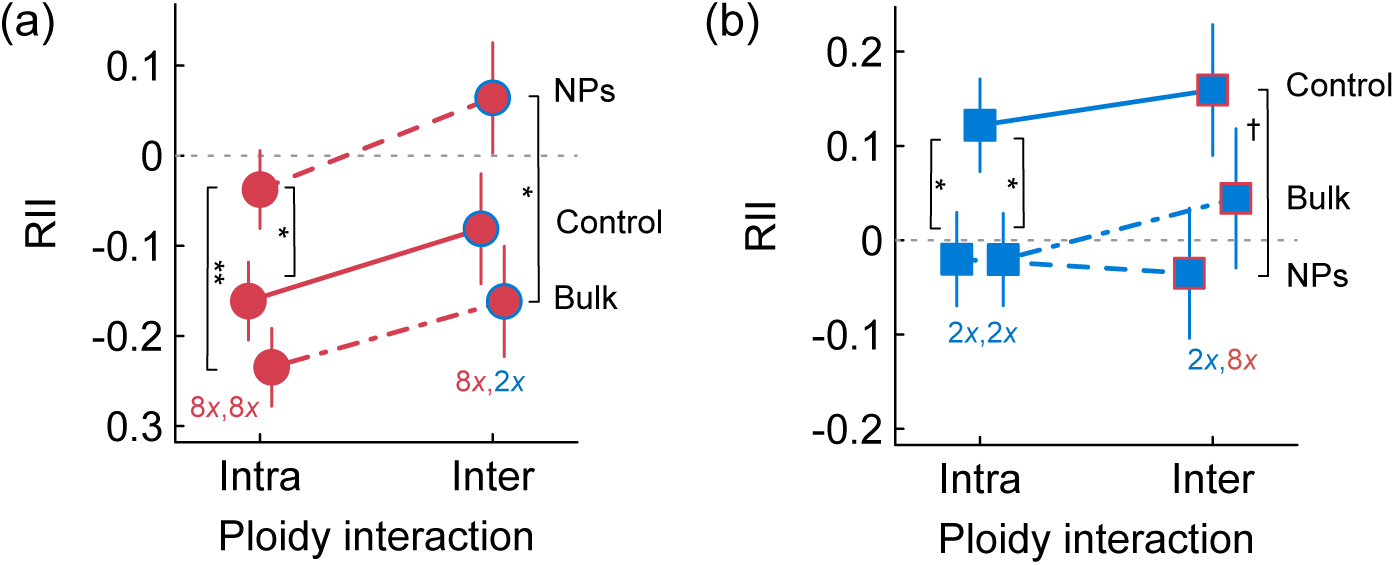
Responses of intraploidy and interploidy interactions to copper oxide nanoparticles (NPs) and bulk particles (Bulk). (a, b) The least-squares mean (LS mean) and 1 SE of the relative interaction index (RII) are plotted for (a) polyploids and (b) diploids estimated from general linear models. Significant contrasts in LS means are denoted: \*\**p* < 0.01; \**p* < 0.05; ^†^*p* = 0.055.

Unlike polyploids, diploids experienced facilitation instead of competition when growing with diploids (RII_2*x*,2*x*_) or polyploids (RII_2*x*,8*x*_) under the control (LS mean: RII_2*x*,2*x*_ *=* 0.12, 95% CI = 0.02–0.22; RII_2*x*,8*x*_ *=* 0.16, 95% CI = 0.02–0.30; Fig. 3b and Fig. S3). These plant–plant interactions became neutral under the NPs and Bulk treatments (Fig. 3b), especially for intraploidy interactions (RII_2*x*,2*x*_; LS mean contrast: control vs. NPs, *F* = 4.2, df = 1, *p* = 0.046; control vs. Bulk, *F* = 4.3, df = 1, *p* = 0.046).

In examining the competitive asymmetry between polyploids and diploids, we found no strong evidence for the hypothesis of polyploid competitive dominance (RII_8*x*,8*x*_ > RII_8*x*,2*x*_ and RII_2*x*,2*x*_ < RII_2*x*,8*x*_; Fig. 1). In polyploids, the overall trend followed the hypothesis (RII_8*x*,8*x*_ > RII_8*x*,2*x*_), but the contrasts of means were not statistically significant across treatments (all *p* > 0.05; Fig. 3a). In diploids, the interactions shifted from facilitation to neutrality (Fig. 3b), different from the proposed competition hypotheses (Figs. S1 and S2).

## 4. Discussion

This study provides, to our knowledge, the first demonstration of how nanoparticles influence species interactions between polyploid and diploid plants. By quantifying intraploidy and interploidy interactions, our results revealed reduced competition in the polyploid under nanoparticle exposure, and a shift towards facilitation when growing with the diploid. This reduction in competitive interactions in the polyploid was primarily caused by nanoscale effects. In contrast, the diploid shifted from facilitation to neutrality under nanoparticle exposure, with a similar reduction in facilitation observed with both nanoparticles and bulk particles. Such contrasting interaction dynamics in the polyploid (reduced competition) and diploid (facilitation to neutrality) translated into observable differences in plant growth (Fig. S3).

### 4.1 Intraploidy and interploidy competitions in the polyploid weakened under nanoparticles

The overall trend of intraploidy and interploidy interactions in the polyploid followed the prediction that polyploids are limited more by individuals of their own than by diploids (Fig. 1 and Fig. S3), with the mean difference not reaching statistical significance due to variation among microcosms. This finding is consistent with observations in other polypoid species complex, including the autotetraploid perennial grass *Dactylis glomerata* under field conditions (Maceira et al., 1993), allotetraploid annual grass *Brachypodium hybridum* in a dry field locality (Rey et al., 2017), and allotetraploid perennial herbaceous *Chrysanthemum indicum* under both well-watering and reduced watering conditions (Guo et al., 2023). However, in the autotetraploid perennial herbaceous *Chamerion angustifolium* (Thompson et al., 2015), similar intraploidy and interploidy competitions were observed under well-watering and reduced watering conditions. Notably, different from the previous studies that involved higher competition densities, such as two to eight plants per pot (Guo et al., 2023), six plants in different pot sizes (Rey et al., 2017), or 16 or 36 plants per pot (Maceira et al., 1993), our microcosm experiment involved competing two plants of the same or different ploidy levels. As competition is density dependent (Zhang and Tielborger, 2020; Guo et al., 2023), the intraploidy and interploidy competitions in the polyploid are expected to intensify with increasing competing individuals, leading to much stronger intraploidy than interploidy competitions than observed here.

Consistent with the stress gradient hypothesis, intraploidy and interploidy competitions in the polyploid were weakened under nanoparticle stress, with a shift towards facilitation when growing with the diploid. This pattern aligns with observations from a limited number of studies investigating the impact of nanoparticles on species interactions (Peng et al., 2017; Wu et al., 2019; Menicagli et al., 2023; Zuo et al., 2024), where a reduction in competition has also been noted under nanoparticle exposure. For example, competitions between species of freshwater green micro-algae were found to decrease under CuO nanoparticles, contributing to species coexistence (Zuo et al., 2024). In ciliate protozoans, the interaction between *Tetrahymena pyriformis* and *Loxocephalus* sp. shifted from competition to facilitation of *Loxocephalus* under cerium oxide (CeO_2_) nanoparticles (Peng et al., 2017). In plants, the competitive effect of the invasive annual herbaceous *Amaranthus retroflexus* on the native *Amaranthus tricolor* decreased under silver (Ag) nanoparticles (Wu et al., 2019). Similarly, the competitive effect of the native perennial grass *Thinopyrum junceum* on the invasive perennial creeping succulent *Carpobrotus* sp. in a coastal dune ecosystem decreased under titanium dioxide (TiO_2_) nanoparticles (Menicagli et al., 2023). The wide range of phylogenetic lineages and diverse nanoparticle types in these previous studies, including our own, underscore the commonality of nanoparticles in reducing species competitions across kingdoms. Future research is needed to discern the factors determining changes in the magnitude (reduced competition) and nature (from competition to facilitation) of species interactions under nanoparticle exposure, such as increased nanoparticle aggregation and changes in nanoparticle surface properties caused by neighboring individuals.

The reduction in competition observed under CuO nanoparticles, but not bulk particles, suggests that the impact was attributable to nanoscale effects. The only other study that investigated the impact of CuO nanoparticles on species competitions in freshwater green micro-algae (Zuo et al., 2024) nevertheless did not include bulk particles to distinguish nanoscale effects vs. material effects. In line with our findings, stronger effects on species interactions under nanoparticles compared to bulk particles were observed in the ciliate protozoans under CeO_2_ nanoparticles (Peng et al., 2017) and in the herbaceous *Amaranthus* plants under Ag nanoparticles (Wu et al., 2019). In addition to species interactions, stronger effects of CuO nanoparticles on individual organisms compared to CuO bulk particles have been found previously. For example, the aquatic plant *Lemna minor* exhibited a significant decrease in fresh biomass within 96 hours of exposure to 10 mg/L CuO NPs, whereas a significant decrease occurred only under exposure to 150 mg/L CuO bulk particles (Song et al., 2016). Likewise, bok choy (*Brassica rapa*) exhibited a significant decrease in leaf dry weight and foliar area under exposure to 75 mg/L CuO NPs when grown in soil for 70 days, whereas a comparable decrease occurred only under exposure to 150 to 300 mg/L CuO bulk particles (Deng et al., 2020). However, exceptions exist; for example, maize (*Zea mays*) seedlings experienced a more pronounced reduction in shoot height and higher root cell membrane injury at 8 mM CuO bulk particles compared to the same concentration of CuO NPs when grown in water for 6 days (Roy et al., 2022), due to higher copper accumulation in seedling roots under CuO bulk particles and possible aggregation reducing the negative impacts of CuO NPs. In our study, we found that the polyploid, when grown in isolation, tended to be more sensitive to CuO NPs than the diploid (Fig. S3). Thus, the presence of neighbors, particularly the less sensitive diploid, likely mitigated nanoparticle stress, resulting in reduced competition for the polyploid and a shift towards facilitation when growing with the diploid. Interestingly, in contrast to the negative impact of CuO NPs, CuO bulk particles tended to benefit the plants when grown in isolation (Fig. S3). As a result, the intraploidy and interploidy competitions in the polyploid were not reduced by CuO bulk particles. Overall, the variability in outcomes from exposure to CuO NPs and bulk particles across studies suggests a context dependency of impacts, influenced by factors such as particle concentrations (low vs. high dosages), environmental complexities (aquatic vs. soil matrix), experimental durations (short vs. long term), interaction types (individual organisms vs. species interactions), species characteristics (e.g., wild vs. cultivated species), and exposure history (prior vs. naïve exposure).

### 4.2 Intraploidy and interploidy interactions shifted from facilitation to neutrality in the diploid under nanoparticles

In this study, we found that the diploid grew better with another diploid or polyploid plant compared to growing alone under control conditions. While such facilitation was unexpected, it suggests potential benefits derived from growing alongside neighbors, such as improved moisture retention in our microcosm experiment. The closed-system microcosms experienced substantial evaporation. As the diploid appeared to grow shorter roots than the polyploid in micropropagation for the axenic microcosms, they were more likely to absorb water from shallower depths, which were more prone to severe evaporation. Growing with a neighbor might have facilitated the upward movement of water into the root zone of the diploid, promoting facilitation. Additionally, growing with another plant might have aided in surface covering, reducing water evaporation. Facilitative effects on neighboring species by enhancing soil moisture retention have been observed in various contexts, such as in the grass *Stipa tenacissima* (Maestre and Cortina, 2004) and *Aristida stricta* (Iacona et al., 2012). Furthermore, growing with neighbors might be more crucial in axenic conditions, as plants lack essential microbial partners, such as mycorrhizal fungi, for improved water access.

In the diploid, intraploidy and interploidy interactions shifted from facilitation to neutrality under CuO NPs and bulk particles. While the diploid was not sensitive to CuO NPs when grown in isolation (Fig. S3), growing with another plant that might help with moisture retention could potentially inadvertently facilitate the release of harmful copper ions and reactive oxygen species, leading to a reduction of facilitative effects compared to control conditions. In contrast, CuO bulk particles tended to benefit the plants when grown in isolation (Fig. S3). Thus, the benefits of growing with neighbors for reasons such as enhanced moisture retention were possibly offset by the drawbacks associated with potential competition from larger plants enhanced by CuO bulk particles. The distinct interaction dynamics of the diploid relative to the polyploid under CuO NPs and bulk particles might be explained by differences in sensitivity between the diploid and polyploid as well as dosage effects. Ploidy difference in sensitivity to copper stress has been observed previously. For example, in the annual herbaceous *Matricaria chamomilla*, diploids were more sensitive to copper stress than colchicine-induced autotetraploids within seven days of exposure to 60 µM of CuCl_2_ (Kováčik et al., 2011) and within 24 hours exposure of 120 µM of CuCl_2_ (Kovacik et al., 2010) when grown hydroponically. Similarly, the diploid *Arabidopsis thaliana* was less tolerant to copper stress than its autotetraploid lines within 14 days of exposure to 50 µM and 100 µM of CuCl_2_ when grown axenically in petri dishes (Li et al., 2017). Such ploidy difference in tolerance to copper stress was often caused by higher accumulation of copper in tissues in diploids than polyploids (Kovacik et al., 2010; Kováčik et al., 2011; Li et al., 2017). Different from the previous studies, in this study, we found that the diploid was not sensitive to CuO NPs, unlike the polyploid (Fig. S3), likely due to its shorter shoots with less surface area for interactions with CuO NPs. While using a widely accepted experimental dosage of 100 ppm CuO NPs and bulk particles, which has been shown to exhibit toxic effects on different plant species (Khan et al., 2021; Gakis et al., 2023), it is important to consider the potential for increased stress in the diploid at higher concentrations of CuO NPs. Furthermore, increased concentrations of CuO bulk particles may diminish the benefits observed in the diploid and polyploid, thereby altering the intricate dynamics of their interactions. Nevertheless, the mechanistic understanding of ploidy-specific responses to nanoparticle stress, their dosage dependency, and the resulting impacts on interactions between diploids and polyploids necessitates future in-depth investigations.

## 5. Conclusions

Our results revealed the impact of nanoparticles on the dynamics of species interactions within a polyploid species complex. Through an axenic experiment, our study directly assessed the influence of nanoparticles on plant–plant interactions, avoiding potential confounding impact from environmental or symbiotic microbes, whose interactions with nanoparticles (Ameen et al., 2021; Gakis et al., 2023; Chen et al., 2024) could otherwise complicate nanoparticle influence. Yet, future studies aiming to quantify both direct and indirect impacts of nanoparticles on species interactions should consider axenic and microbe-addition comparisons. However, due to the need of microcosms for axenic manipulation, our study has limitations. New investigations are, therefore, needed to provide insights into: (1) how nanoparticle effects on species interactions vary with competitor density and exposure intensity, (2) the short-term versus long-term impacts of nanoparticles, and (3) the extent to which findings from controlled experiments can be extrapolated to field performance. Moreover, our study focused on established polyploids and diploids. Thus, the intraploidy and interploidy interactions may reflect the combined outcome of whole-genome duplication and polyploidy-enabled adaptation in response to nanoparticles. To disentangle the effect of whole-genome duplication in mediating species interactions under nanoparticle exposure, similar tests in natural or synthetic neopolyploids across polyploid species complex and genotypes are needed in both wild and domesticated plants, in face of growing nanotechnology in agriculture and other industries (Lowry et al., 2019; Keller et al., 2023). These studies together will advance our understanding of the potential off target impacts of nanoparticles on organisms and communities, offering critical insights for environmental and organismal conservation efforts.

## Acknowledgements

We thank Charlotte Hewins for the assistance in plant cultivation and Jessica LaBella for the assistance in experimental setup. We also thank Tia-Lynn Ashman and Aaron Liston for providing the wild strawberry seeds.

## Appendix A. Supplementary material

Supplementary data associated with this article can be found in the online version.

## Data Availability

Data and analyses that support the findings of this study are included in this published article and its electronic supplementary material and on Mendeley Data (DOI: 10.17632/w6wkf647s3.1).

## CRediT authorship contribution statement

**Elizabeth A. Esser:** Investigation, Writing - Original Draft, Writing-review and editing, Visualization. **Jiaqi Tan:** Resources, Writing - Review & Editing. **Na Wei:** Conceptualization, Methodology, Formal analysis, Resources, Writing - Original Draft, Writing-review and editing, Visualization, Supervision, Project administration.

## Supplementary Material

**Fig. S1.**
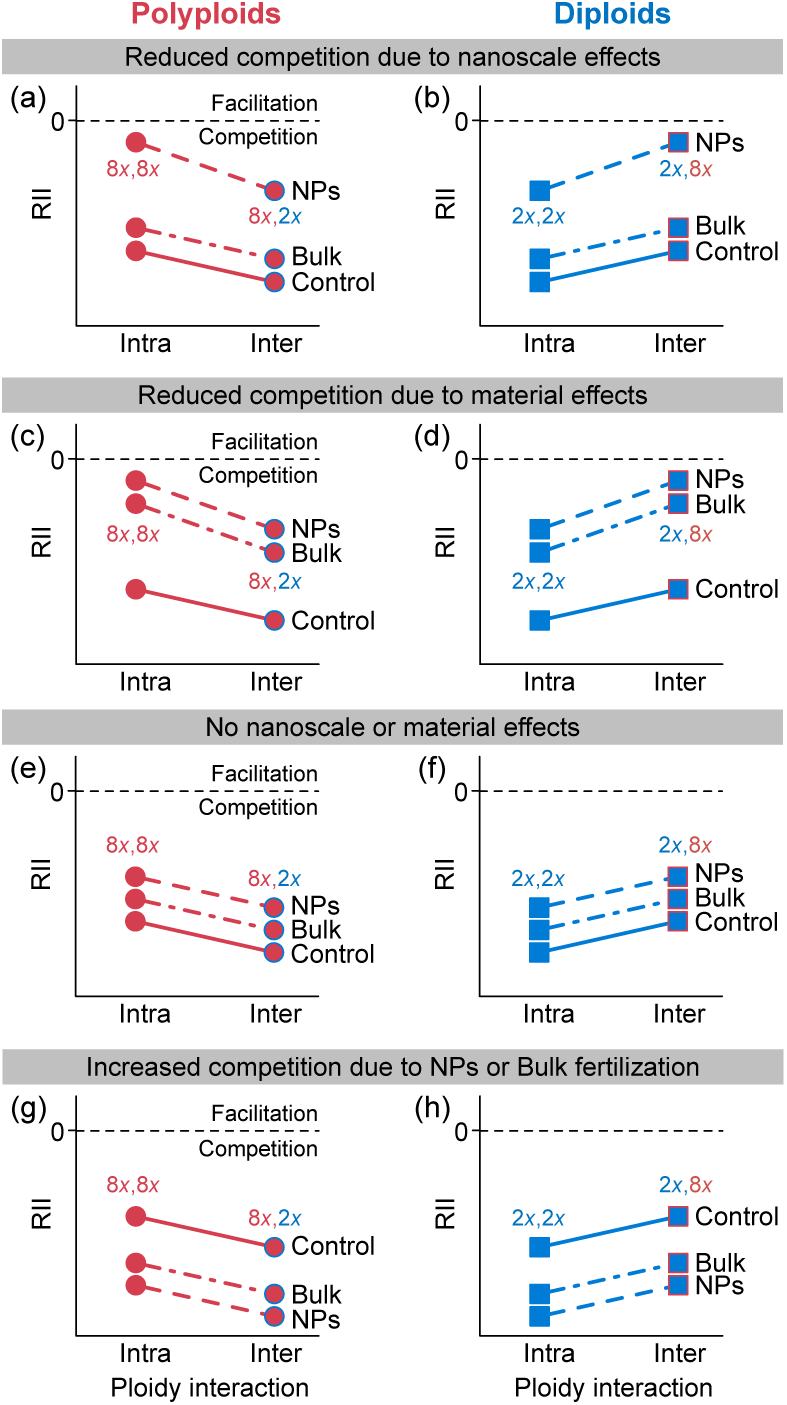
Hypotheses of competitive interactions under diploid dominance in response to nanoparticles. (a-f) Plant–plant interactions between ploidy levels are quantified using the relative interaction index (RII), where RII < 0 indicates competition and RII > 0 indicates facilitation. If diploids are stronger competitors, the competitive effect of diploids on polyploids (8*x*, octoploid as an example; RII_8*x*,2*x*_) is expected to be stronger than the effect of polyploids on polyploids (RII_8*x*,8*x*_), resulting in stronger interploidy than intraploidy competitions, RII_8*x*,2*x*_ > RII_8*x*,8*x*_. In contrast, diploids are expected to be more affected by intraploidy competition (RII_2*x*,2*x*_) than interploidy competition (RII_2*x*,8*x*_), leading to RII_2*x*,2*x*_ > RII_2*x*,8*x*_. The intraploidy and interploidy competitions may, nevertheless, change under non-resource stress (e.g., nanoparticles and bulk particles of the material). The stress gradient hypothesis (SGH) predicts reduced competition with increased stress. (a, b) If the stress is caused by nanoscale effects rather than material effects, a reduction in competition is expected under nanoparticles (NPs), in contrast to bulk particles (Bulk) and the control (Control) where no nanoparticles or bulk particles are introduced. (c, d) If the stress is caused by material effects, a reduction in competition is expected to be similar under both nanoparticles and bulk particles, relative to the control. (e, f) However, competition may remain unchanged if nanoparticles and bulk particles are not the primary sources of stress that influences plant–plant interactions. (g, h) On the other hand, if nanoparticles and bulk particles provide fertilization, intraploidy and interploidy competitions may increase due to reasons such as increased plant sizes.

**Fig. S2.**
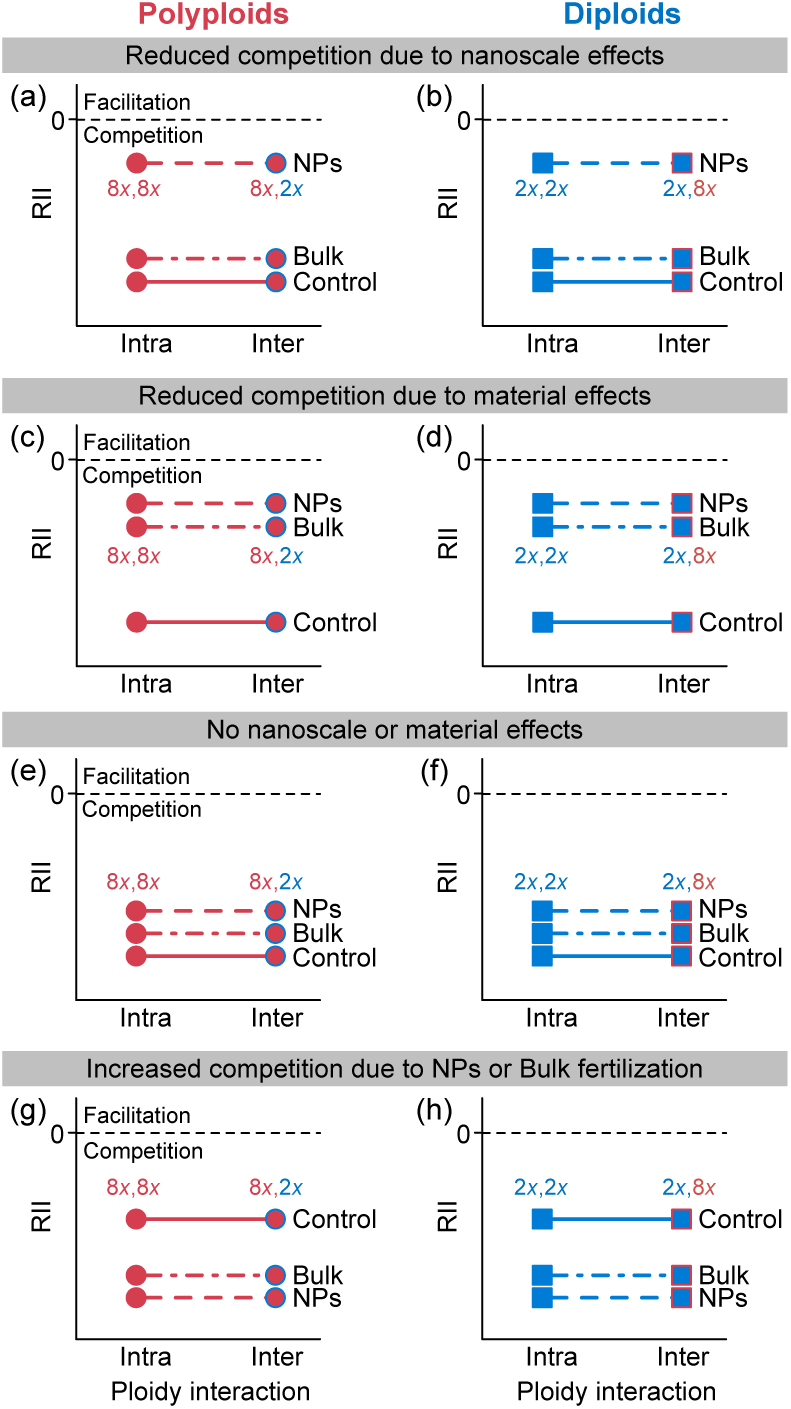
Hypotheses of interactions under competitive symmetry between polyploids and diploids in response to nanoparticles. (a-f) Plant–plant interactions between ploidy levels are quantified using the relative interaction index (RII), where RII < 0 indicates competition and RII > 0 indicates facilitation. In the scenario of competitive symmetry between polyploids and diploids, intraploidy and interploidy competitions are expected to be similar: RII_8*x*,2*x*_ = RII_8*x*,8*x*_ and RII_2*x*,2*x*_ = RII_2*x*,8*x*_. The intraploidy and interploidy competitions may, nevertheless, change under non-resource stress (e.g., nanoparticles and bulk particles of the material). The stress gradient hypothesis (SGH) predicts reduced competition with increased stress. (a, b) If the stress is caused by nanoscale effects rather than material effects, a reduction in competition is expected under nanoparticles (NPs), in contrast to bulk particles (Bulk) and the control (Control) where no nanoparticles or bulk particles are introduced. (c, d) If the stress is caused by material effects, a reduction in competition is expected to be similar under both nanoparticles and bulk particles, relative to the control. (e, f) However, competition may remain unchanged if nanoparticles and bulk particles are not the primary sources of stress that influences plant–plant interactions. (g, h) On the other hand, if nanoparticles and bulk particles provide fertilization, intraploidy and interploidy competitions may increase due to reasons such as increased plant sizes.

**Fig. S3.**
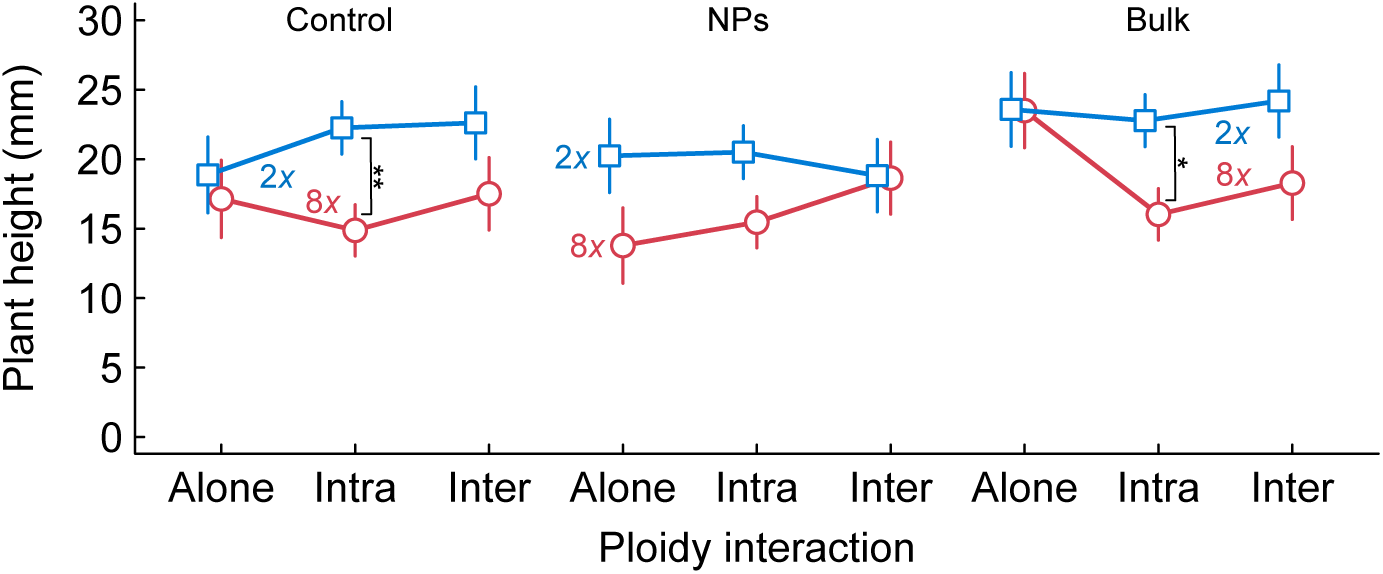
Plant height under copper oxide nanoparticles (NPs, 100 ppm) and bulk particles (Bulk, 100 ppm) and the control. The least-squares mean (LS mean) and 1 SE are plotted for polyploids (8*x*, open circles) and diploids (2*x*, open squares) estimated from a general linear model with predictors of ploidy (8*x* and 2*x*), treatment (Control, NPs, and Bulk), and competition (growing alone, intraploidy, and interploidy), as well as their two-way and three-way interactions, with the initial plant size as a covariate. Significant contrasts in LS means are denoted: \*\**p* < 0.01; \**p* < 0.05.

**Table S1.**
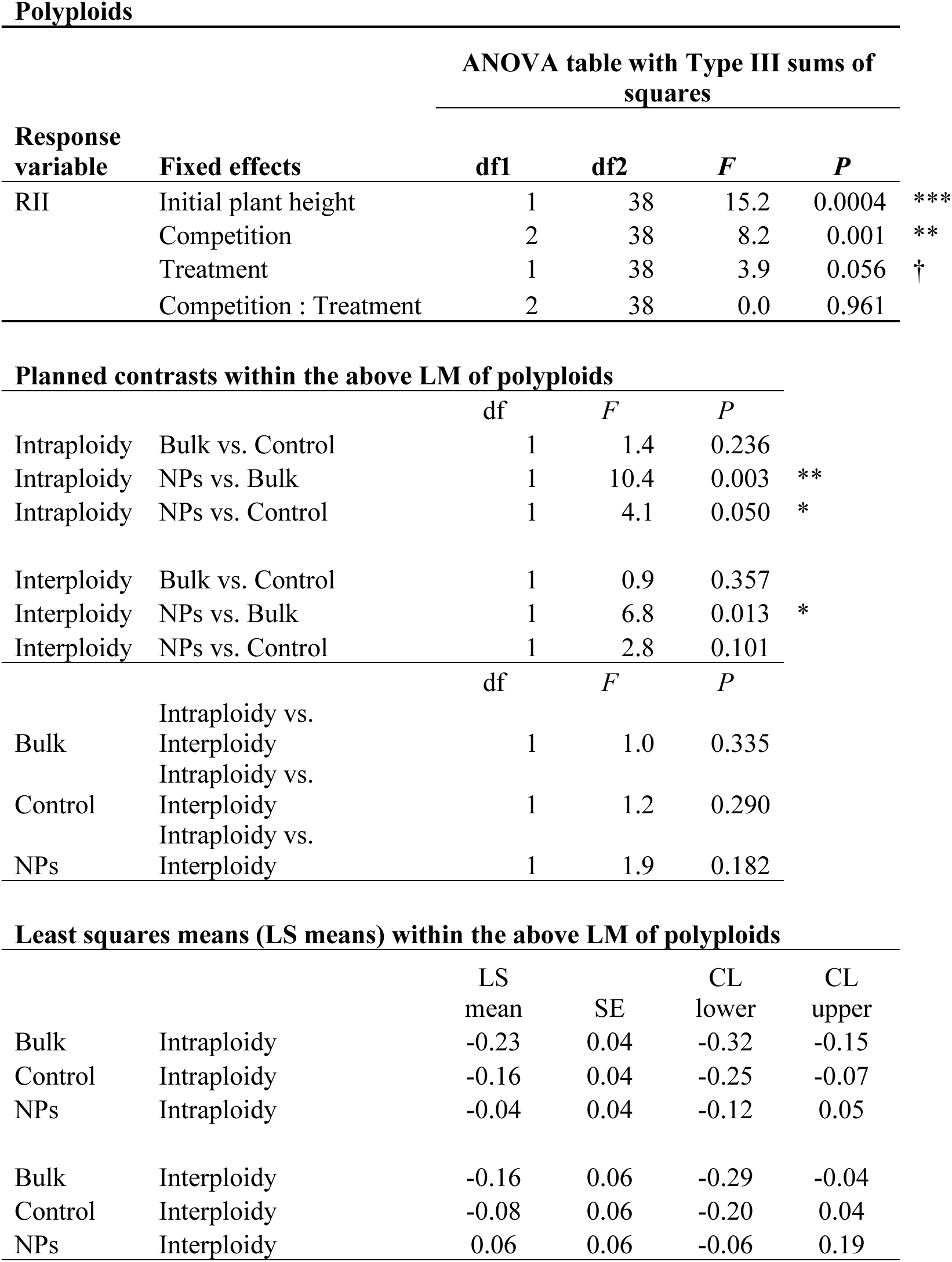
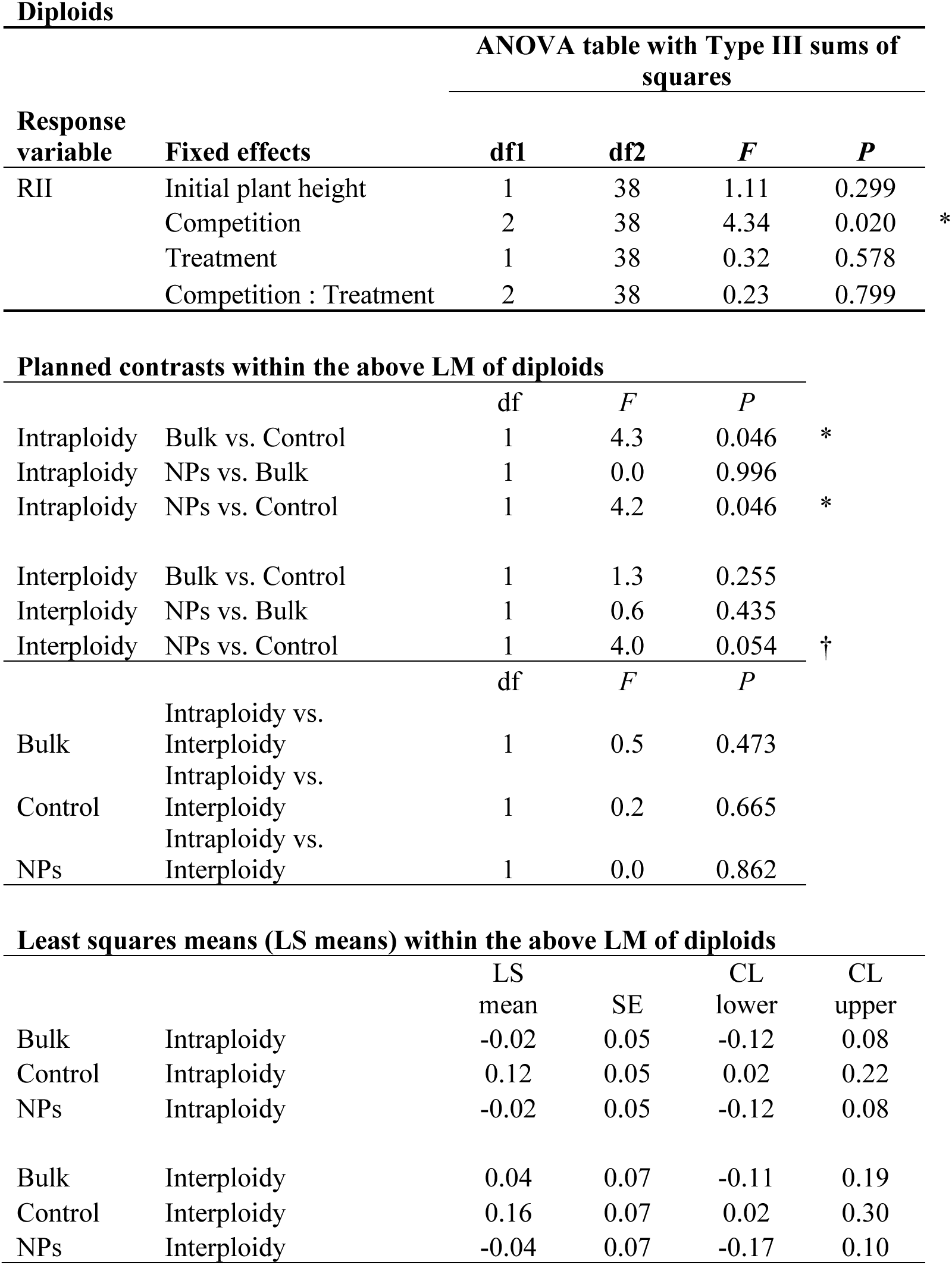
Models and planned contrasts examining intraploidy and interploidy interactions in polyploids and diploids. Linear model (LM) specification: Predictors: Initial plant height + Competition (Intraploidy, Interploidy) + Treatment (NPs, Bulk, Control) + Competition : Treatment

